# On reconstruction of cortical functional maps using subject-specific geometric and connectome eigenmodes

**DOI:** 10.1101/2024.10.28.620635

**Authors:** Anders S. Olsen, Sina Mansour L., James C. Pang, Andrew Zalesky, Dimitri Van De Ville, Hamid Behjat

**Affiliations:** Department of Applied Mathematics and Computer Science, Technical University of Denmark, Kgs. Lyngby, Denmark; Centre for Sleep & Cognition & Centre for Translational Magnetic Resonance Research, National University of Singapore, Singapore; Melbourne Neuropsychiatry Centre, Department of Psychiatry, The University of Melbourne and Melbourne Health, Victoria, Australia; School of Psychological Sciences, Turner Institute for Brain and Mental Health, Monash University, Victoria, Australia; Department of Psychiatry, Melbourne Medical School, The University of Melbourne, Melbourne, Australia; Neuro-X Institute, École Polytechnique Fédérale de Lausanne, Geneva, Switzerland; Department of Radiology and Medical Informatics, University of Geneva, Geneva, Switzerland; Department of Clinical Sciences Malmö, Faculty of Medicine, Lund University, Lund, Sweden

**Keywords:** cortical geometry, structural connectivity, functional MRI, brain structure-function interplay

## Abstract

Understanding the interplay between human brain structure and function is crucial to discern neural dynamics. This study explores the relation between brain structure and macroscale functional activity using subject-specific structural connectome eigenmodes, complementing prior work that focused on group-level models and geometry. Leveraging data from the Human Connectome Project, we assess accuracy in reconstructing various functional MRI-based cortical maps using individualised eigenmodes, specifically, across a range of connectome construction parameters. Our results show only minor differences in performance between surface geometric eigenmodes, a local neighborhood graph, a highly smoothed null model, and individual and group-level connectomes at modest smoothing and density levels. Furthermore, our results suggest that spatially smooth eigenmodes best explain functional data. The absence of improvement of individual connectomes and surface geometry over smoothed null models calls for further methodological innovation to better quantify and understand the degree to which brain structure constrains brain function.

## 1. INTRODUCTION

Orchestrated regional and distributed activity constitutes a fundamental aspect of neural dynamics. Integration of cross-regional activity involves signal propagation along axonal fibers as well as traversal along the cortical sheet, adhering to the principles of neural field theory [1]. In a recent study [2], the relation between the brain’s geometry, i.e., cortical folding pattern, and functional activity—encompassing a range of subject-specific task maps and resting-state functional MRI signals—was scrutinized using eigenmodes of the Laplace-Beltrami operator of a cortical surface mesh. The findings suggested that surface geometric eigenmodes offered a more parsimonious description of brain functional maps than eigenmodes of a structural connectome. A subsequent work [3] explored a wider range of parameters associated to connectome construction (connectome density, spatial smoothing [4], and weight binarization) and found that by using appropriate refinements, connectome eigenmodes could achieve reconstruction accuracies comparable to those of geometric eigenmodes. Importantly, these prior work focused on *group-level* geometric and connectome representations, thus, overlooking subtle subject-specific idiosyncrasies that are inherent in the relationship between structure and function [5]–[7].

In this study, we hypothesized that eigenmodes of subject-specific connectomes would better reconstruct subject-specific functional maps than those of group-level connectome and surface geometry. We investigated the performance at various connectome construction parameters and benchmarked our results against a null model that consisted of smoothed eigenmodes derived from noise. Following previous studies, we reconstructed lightly smoothed task maps in template space, but also unsmoothed stationary co-activation pattern (CAP) maps computed in native space [8] to avoid any implicit smoothing due to nonlinear transformation to template space. All processing and analysis scripts used in this study are made available^1^.

## 2. METHODS

### 2.1. Data

We used anatomical, diffusion, and functional MRI data from the Human Connectome Project (HCP) [9], specifically the 100 unrelated subjects set, study complied with ethical standards. As in previous studies [2], we focused exclusively on the left hemisphere in HCP’s fs-LR 32k surface template comprising *N* = 29, 696 vertices after excluding the medial wall. We evaluated 47 task contrast maps, where each map was processed using HCP’s protocol that involves nonlinear projection into MNI space and surface smoothing with a 2mm FWHM Gaussian kernel [10]. Furthermore, we used stationary co-activation pattern (CAP), a proxy for the seed-based functional connectivity map, as constructed in [8] from the left-right phase encoding resting-state fMRI data (REST 1) of each individual; one volumetric map per cortical parcel of the Schaefer-400 atlas [11]. Each map was projected onto the native pial surface (∼ 130K vertices) and subsequently resampled onto the 32k HCP-resolution using nearest-neighbor vertex matching.

### 2.2. Brain structure: comparing geometry and connectome

We study two approaches for representing brain structure: cortical surface geometry and high-resolution structural connectivity; see first column in Fig. 1. The geometry of the cerebral cortex can be represented as a mesh that encodes geodesic adjacency of cortical nodes given their convoluted foldings as well as their physical location in space, which, in turn, allows the calculation of cortical curvature. Here we investigated subject-specific and template mid-thickness surface files [12], which are calculated as the midpoint between the pial and white matter surfaces provided by the FreeSurfer software suite [13]. In contrast, a structural connectome encodes any potential white matter-mediated local and distal structural connection between cortical regions. The presence or absence of connections and their strength are determined by counting the number of tractography streamlines in an adjacency matrix, however, without encoding the spatial location of nodes. We performed tractography using MRTrix3 [14], confined to 20 million streamlines in total per subject; further details can be found in [15]. For the purpose of creating a null model, we constructed a matrix of size *N* × 200 composed of random univariate Gaussian noise elements; we used 200 columns to obtain a null basis set with the same cardinality as that of geometry and connectome.

**Fig. 1.**
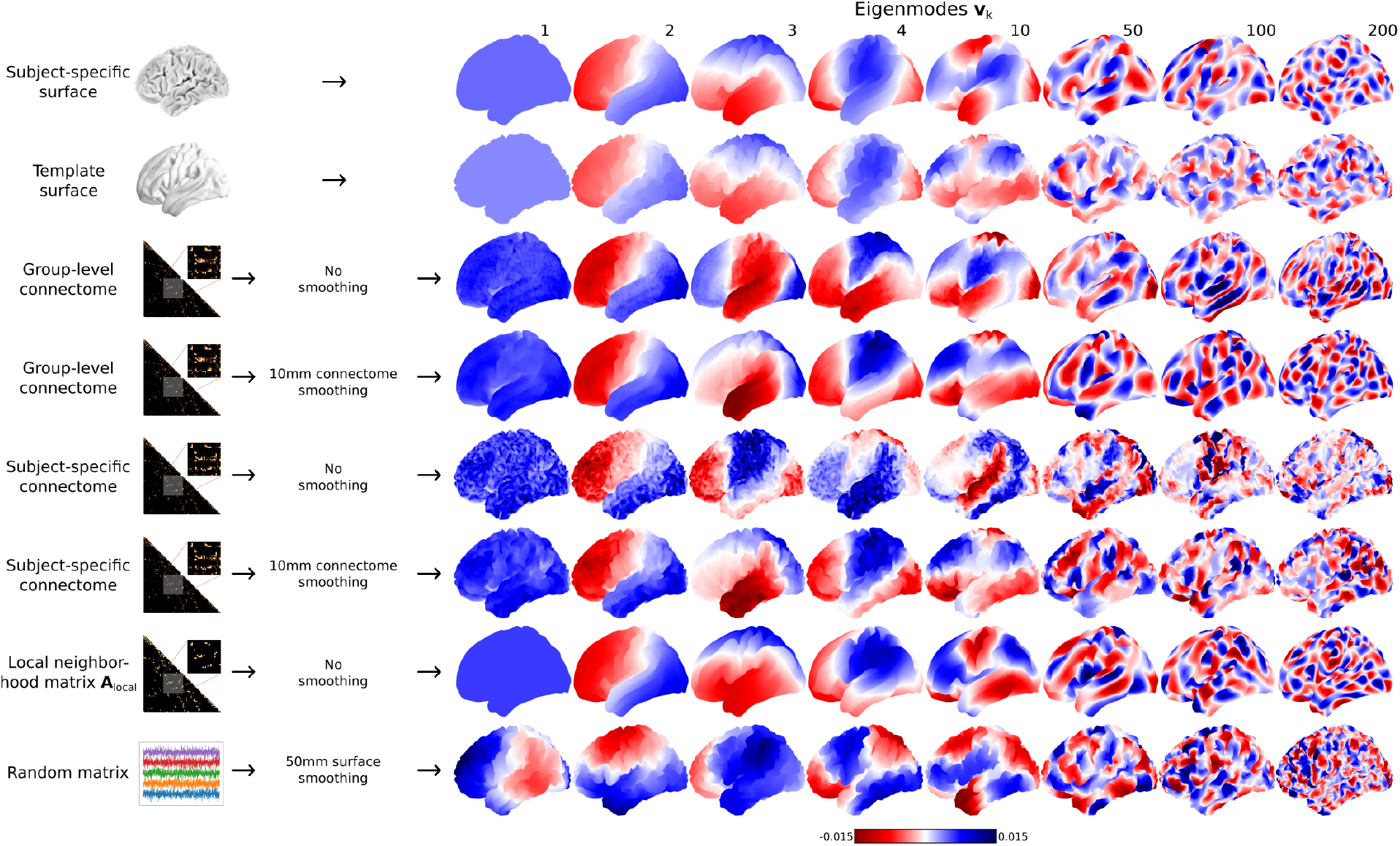
Brain eigenmodes derived from surface, null model, or connectome representations of brain structure across different settings.

### 2.3. Surface and connectome spatial smoothing

It is important to differentiate between two types of smoothing operations used in this work: *connectome spatial smoothing* [4] and *surface smoothing*; see second column in Fig. 1. For connectomes, we used connectome spatial smoothing, which smooths streamline counts across neighboring nodes as defined by the subject’s surface mesh. We tested for smoothing kernel sizes of 0, 2, 6, and 10mm FWHM, and all smoothed connectomes were subsequently thresholded at an absolute value of 0.1 to alleviate storage issues. For the null model, the random matrix was surface smoothed; we used the connectome workbench^2^, wherein the physical distances between nodes encoded in the mid-thickness surface file were used. We investigated changes in reconstruction accuracy as a function of smoothing kernel sizes from 0 (no smoothing) to 70mm FWHM.

### 2.4. Connectome, geometric, and null model eigenmodes

We constructed group-level connectomes by averaging individual connectomes across the 100 subjects. By applying connectome spatial smoothing (four smoothing levels: 0, 2, 6, 10 mm FWHM) and subsequent thresholding (five density levels: 10^−5^, 10^−4^, 10^−3^, 0.01, and 0.05), we obtained 20 connectomes per subject and 20 connectomes at the group-level. Each connectome can be represented as a weighted graph adjacency matrix **W** of size *N* × *N*. To ensure graph connectedness, a local neighborhood matrix **A**_local_, containing ones for the direct neighbors of each vertex and zeros otherwise, was added: **A** = **W** + **A**_local_. In the following, we used **A** in both weighted and binary form.

For each **A**, the normalized graph Laplacian was computed **L** = **I** − **D**^−1*/*2^**AD**^−1*/*2^, where **D** denotes the diagonal degree matrix and **I** denotes the identity matrix. **L** was then eigendecomposed, **L** = **VΛV**^⊤^, where **V** is the matrix of eigenvectors 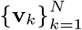 and **Λ** is the diagonal matrix of eigenvalues. The Laplacian eigenvectors (a.k.a. *eigenmodes*) are ordered according to low-to-high eigenvalues representing a proxy for spatial frequency. As in prior work [2], [3], we computed and studied only the first 200 eigenmodes. Surface geometric eigenmodes were computed via the eigendecomposition of the Laplace-Beltrami operator using the LaPy library [16], [17]. Eigenmodes of surface-smoothed null model were computed using the singular value decomposition, where modes were sorted according to decreasing singular value. Surface projections of a representative set of eigenmodes across various settings investigated in this study are shown on the right-hand side of Fig. 1.

### 2.5. Reconstruction of cortical maps using eigenmodes

Considering a graph signal processing perspective [18], a graph representation of the brain—e.g., cortical surface mesh or structural connectome—can be viewed as a backbone on which brain function resides [19], [20]. In such a setting, the eigenmodes of a suitable shift operator defined on the graph can be leveraged as a basis to decompose brain function, aiming to unravel intricacies of brain structure-function interplay [21]–[23]. To evaluate the efficacy of the derived eigenmodes (cf. Section 2.4) in explaining functional maps, linear model coefficients were computed for each basis by incrementally including eigenmodes, i.e.,

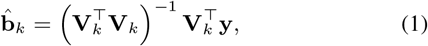

where **y** ∈ ℝ^*N*^ denotes the vectorized form of an fMRI map defined on the cortical surface— a.k.a. a graph signal, see Fig. 2(a)—and **V**_*k*_ = [**v**_1_, **v**_2_, …, **v**_*k*_], where *k* = 1, …, 200. Model coefficients were then used to reconstruct the maps as 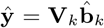. Reconstruction accuracy was computed via the Pearson correlation coefficient between **y** and **ŷ**. To prevent any bias in reconstruction accuracy from smoothing out vertex-level noise, we did not downsample the cortical maps [2], [3] prior to computing the correlations.

**Fig. 2.**
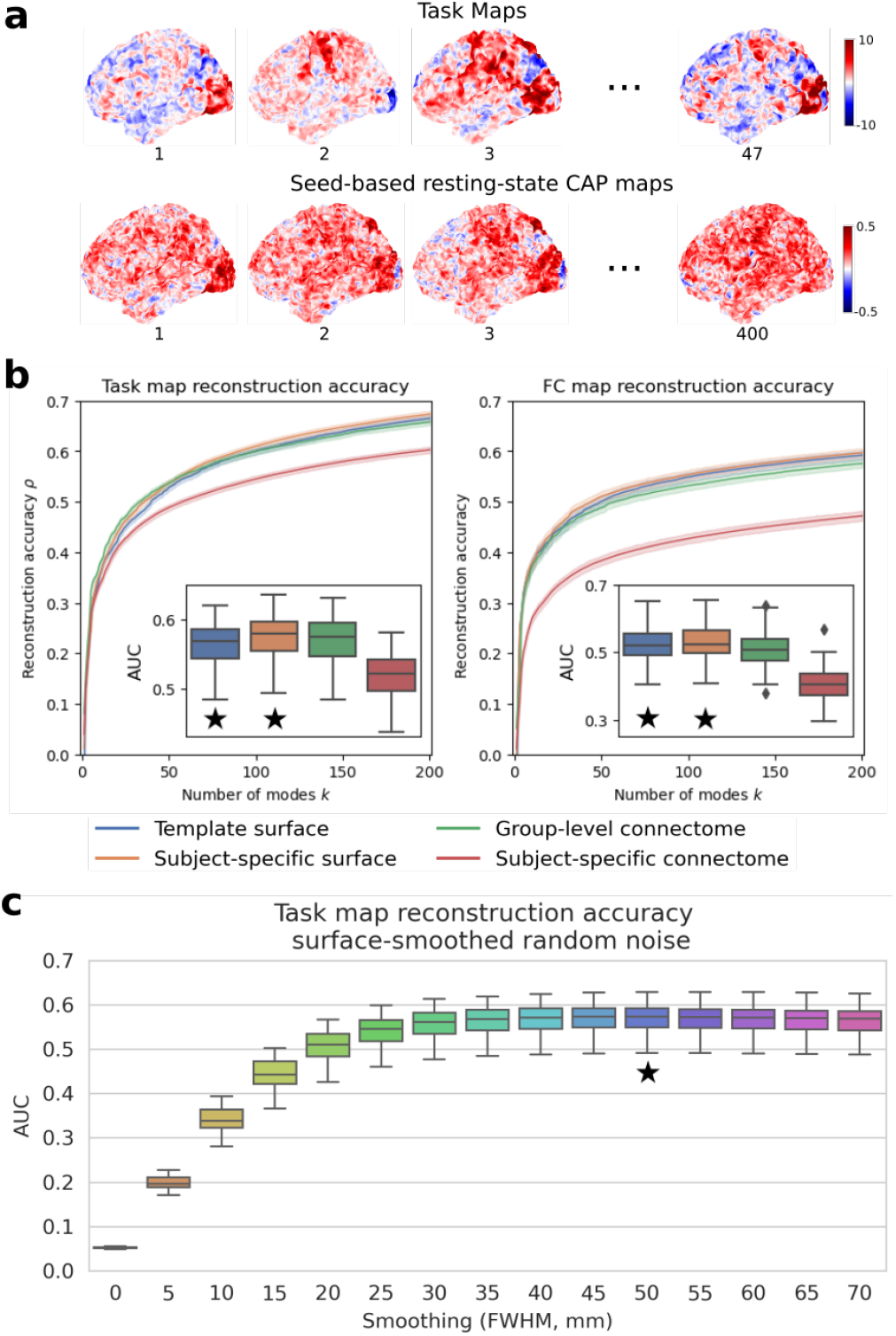
(a) Surface projection of task and resting-state CAP maps of a representative subject. (b) Reconstruction accuracy of cortical maps as a function of the number of included eigenmodes (from high to low spatial wavelengths). Incrementally including more eigenmodes increases accuracy, summarized by the area under the curve (AUC). (c) Efficacy of a spatially smoothed null model eigenmodes in explaining task maps. Box plots in (b) and (c) show distribution across subjects and those that are replicated in Fig. 3 are marked with ⋆.

## 3. RESULTS AND DISCUSSION

Fig. 2(b) compares the reconstruction accuracies between four key settings (template/group-level vs subject-specific) on task and resting-state maps. Eigenmodes of subject-specific connectomes perform worse, whereas those of subject-specific surfaces perform slightly better than the other two settings. Fig. 2(c) shows the reconstruction accuracies using surface-smoothed null model eigenmodes, wherein performance increases with an increased amount of smoothing, reaching a plateau at 50mm FWHM.

Fig. 3 shows the reconstruction accuracies using eigenmodes of geometry, null model, **A**_local_, and connectomes at various parameters. Smoothed, high-density group-level/subject-specific weighted connectomes achieved similar performance as geometric eigen-modes, although no model outperformed **A**_local_ and the null model. Reconstruction accuracies for CAP maps were generally slightly lower than those for task maps, although the relation between parameter settings and surface-based models remained unchanged.

**Fig. 3.**
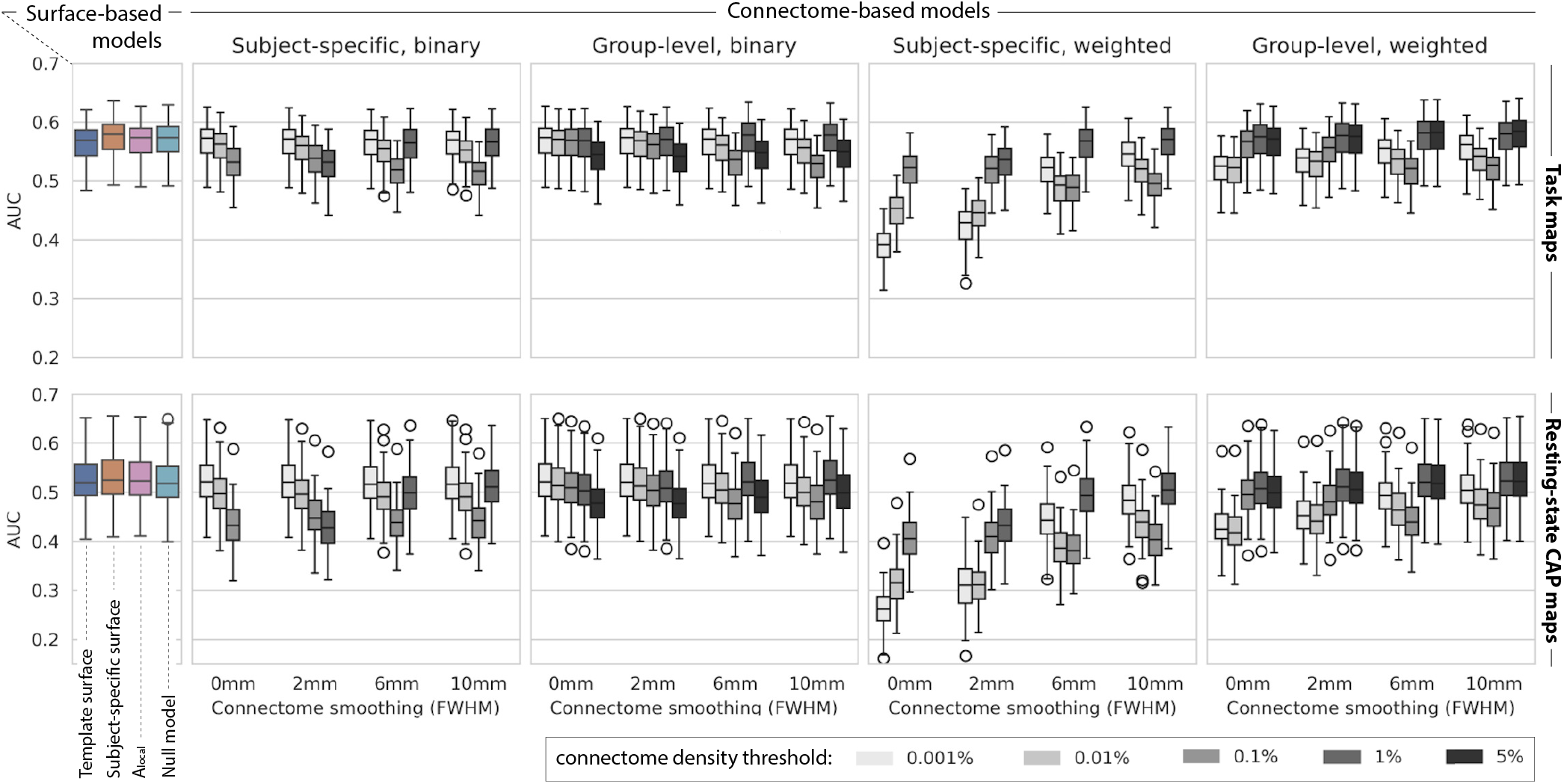
Reconstruction accuracy using surface, connectome, and null model eigenmodes for task (average across 47 tasks) and resting-state CAPs (average across maps associated to 400 brain regions); box plots show distribution of AUC across subjects. For subject-specific connectomes, results for high-density thresholds (5% and/or 1%) are missing since not all connectomes had such high densities.

Reconstruction accuracy using minimally smoothed binary connectomes consistently decreased with higher density, suggesting that binarization reduces reconstruction accuracy for connectomes that entail a larger number of edges. Upon introducing higher connectome density and connectome-spatial-smoothing—which smooths and narrows the distribution of edge weights—this effect is reduced suggesting that modest smoothing and high density are beneficial for connectomes. These results corroborate group-level results in [3]. Binarization results in decreased performance for 5%-density group-level connectomes, suggesting that there is an edge-weight limit, below which binarization results in inflating the importance of spurious edges. Group-level connectomes exhibited slightly increased performance compared to individual connectomes, an effect consistent across tasks and seed-based CAP maps.

Reconstruction using **A**_local_, surprisingly, resulted in similar performance with connectome and geometric eigenmodes. The decrease in performance from low-density to mid-density (0.1%) binary connectomes can be interpreted as a departure from **A**_local_, across smoothing levels. Conversely, for weighted unsmoothed and low-smoothed (2mm) connectomes, low density results in poor performance, which is likely related to included weights having high value while **A**_local_ is binary (cf. definition of **A** under Section 2.4). These results suggest that **A**_local_ itself entails a well-performing eigenbasis, while connectomes with a high level of information, i.e., high density and high smoothing, also perform well; connectomes with parameters in between, including binarization, perform poorly. Unlike surface geometry, and by extension the surface-smoothed null model, **A**_local_ encodes only the immediate three direct neighbors of each vertex, without capturing their spatial location or curvature, implying that brain’s folding patterns, the morphology of which is shown to be akin to a sphere [24], may not be necessary for explaining macro-scale functional patterns, as also suggested in [25].

The eigenmodes of the well-performing bases all appear visually smooth (Fig. 1) relative to the foldings of the cortical surface, and with clear spatial directionality, whereas those of the unsmoothed subject-specific connectome that performed worse are more jagged and individual areas have more influence. These results suggest that eigenmodes that entail smooth, brain-wide coverage are beneficial to effectively reconstruct functional task and CAP maps.

Overall, our results suggest little difference between structural brain models in explaining functional imaging data, whether the models are geometric, smoothed connectomic, or a smoothed random basis. Sorting the eigenmodes according to estimated linear model coefficients 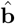 improved overall reconstruction accuracies (results not shown) but did not change the relation between models in terms of reconstruction performance. There are a few limitations in this study that are worth highlighting. Firstly, it is important to note that here we only evaluated the first 200 eigenmodes to align our results with prior work [2], [3]—and in part for computational reasons—whereas results in Fig. 2(b) clearly show that, on average, only 45-70% of variance in functional maps are explained by this subset of eigenmodes, a finding that is in line with other related studies on using brain eigenmodes [23], [26]. Future studies could benefit from exploring a larger range of eigenmodes, for instance evaluating up to a spectral point where 80-90% of the variance in the maps are explained, ideally, coupled with employing novel strategies to study the role of individual eigenmodes (or spectral bands [27]) in characterizing subtle functional spatial patterns rather than using overall reconstruction accuracy as the measure of validation. Secondly, we thresholded our connectomes using an absolute value of 0.1 which might have an effect on connectome eigenmodes. We noticed that well-performing eigenmodes generally showed brainwide coverage and believe this to be related to the “smoothness” of the eigenbasis, however, future studies should further develop this aspect with a quantitative measure of smoothness. The inclusion of **A**_local_ in the connectomes ensures graph connectedness but greatly limits the interpretational separability of tractographic from geometric eigenmodes. Sampling enough streamlines such that the inclusion of **A**_local_ is unnecessary would be beneficial, albeit computationally challenging. Here we sampled 20 million streamlines and observed no change in reconstruction accuracy over 5 million streamlines (not shown). Alternatively, voxel-wise whole brain graphs [23] that entail both gray and white matter may be explored in future work to not only overcome challenges related to tractography in building high-resolution connectomes, but also to integrate functional mediation patterns [28], [29] in white matter when interpreting cortical functional patterns.

## 4. CONCLUSION

When using modest connectome smoothing density, our findings generally suggest little difference in reconstruction accuracy between surface geometry, structural connectomes (both individual and group-level), a smoothed null model, and a local neighborhood graph. We believe that the current framework (total reconstruction accuracy using a set of modes) favors smooth eigenmodes to achieve high graph signal reconstruction accuracies. Future work can benefit from using alternative methods to better quantify the contribution of each eigenmode, or sets of eigenmodes across the spectrum, to illuminate the intricate link between subtleties of brain structure and function at the individual level.

## Footnotes

1 https://github.com/anders-s-olsen/BrainGSP_subject-specific

2 https://www.humanconnectome.org/software/connectome-workbench

## References

[1] V. K. Jirsa and H. Haken, “Field Theory of Electromagnetic Brain Activity,” Physical Review Letters, 1996. DOI: 10.1103/PhysRevLett.77.960.

[2] J. C. Pang, K. M. Aquino, M. Oldehinkel, P. A. Robinson, B. D. Fulcher, M. Breakspear, and A. Fornito, “Geometric constraints on human brain function,” en, Nature, 2023. DOI: 10.1038/s41586-023-06098-1.

[3] S. Mansour L, H. Behjat, D. V. D. Ville, R. E. Smith, B. T. T. Yeo, and A. Zalesky, Eigenmodes of the brain: Revisiting connectomics and geometry, en, 2024. DOI: 10.1101/2024.04.16.589843.

[4] S. Mansour L, C. Seguin, R. E. Smith, and A. Zalesky, “Connectome spatial smoothing (CSS): Concepts, methods, and evaluation,” eng, NeuroImage, 2022. DOI: 10.1016/j.neuroimage.2022.118930.

[5] S. J. Harrison, J. D. Bijsterbosch, A. R. Segerdahl, S. P. Fitzgibbon, S.-R. Farahibozorg, E. P. Duff, S. M. Smith, and M. W. Woolrich, “Modelling subject variability in the spatial and temporal characteristics of functional modes,” NeuroImage, 2020.

[6] J. D. Bijsterbosch, C. F. Beckmann, M. W. Woolrich, S. M. Smith, and S. J. Harrison, “The relationship between spatial configuration and functional connectivity of brain regions revisited,” Elife, 2019.

[7] A. Rajesh, N. A. Seider, D. J. Newbold, B. Adeyemo, S. Marek, D. J. Greene, A. Z. Snyder, J. S. Shimony, T. O. Laumann, N. U. Dosenbach, et al., “Structure–function coupling in highly sampled individual brains,” Cerebral Cortex, 2024.

[8] C. Ferritto, M. G. Preti, S. Moia, D. Van De Ville, and H. Behjat, “Brain Fingerprinting Using FMRI Spectral Signatures On High-Resolution Cortical Graphs,” en, in 2023 IEEE International Conference on Acoustics, Speech, and Signal Processing Workshops (ICASSPW), 2023. DOI: 10.1109/ICASSPW59220.2023.10193247.

[9] D. C. Van Essen, S. M. Smith, D. M. Barch, T. E. Behrens, E. Yacoub, and K. Ugurbil, “The WU-Minn Human Connectome Project: An overview,” NeuroImage, 2013. DOI: 10.1016/J.NEUROIMAGE.2013.05.041.

[10] M. F. Glasser, S. N. Sotiropoulos, J. A. Wilson, T. S. Coalson, B. Fischl, J. L. Andersson, J. Xu, S. Jbabdi, M. Webster, J. R. Polimeni, D. C. Van Essen, and M. Jenkinson, “The minimal preprocessing pipelines for the Human Connectome Project,” NeuroImage, 2013. DOI: 10.1016/j.neuroimage.2013.04.127.

[11] A. Schaefer, R. Kong, E. M. Gordon, T. O. Laumann, X.-N. Zuo, A. J. Holmes, S. B. Eickhoff, and B. T. T. Yeo, “Local-Global Parcel-lation of the Human Cerebral Cortex from Intrinsic Functional Connectivity MRI,” Cerebral Cortex, 2018. DOI: 10.1093/cercor/bhx179.

[12] D. C. Van Essen, M. F. Glasser, D. L. Dierker, J. Harwell, and T. Coalson, “Parcellations and hemispheric asymmetries of human cerebral cortex analyzed on surface-based atlases,” eng, Cerebral Cortex (New York, N.Y.: 1991), 2012. DOI: 10.1093/cercor/bhr291.

[13] B. Fischl, “FreeSurfer,” NeuroImage, 2012. DOI: 10.1016/j.neuroimage.2012.01.021.

[14] J.-D. Tournier, R. Smith, D. Raffelt, R. Tabbara, T. Dhollander, M. Pietsch, D. Christiaens, B. Jeurissen, C.-H. Yeh, and A. Connelly, “MRtrix3: A fast, flexible and open software framework for medical image processing and visualisation,” NeuroImage, 2019. DOI: 10.1016/j.neuroimage.2019.116137.

[15] S. Mansour L, M. A. Di Biase, R. E. Smith, A. Zalesky, and C. Seguin, “Connectomes for 40,000 UK Biobank participants: A multi-modal, multi-scale brain network resource,” eng, NeuroImage, 2023. DOI: 10.1016/j.neuroimage.2023.120407.

[16] M. Reuter, F.-E. Wolter, and N. Peinecke, “Laplace–Beltrami spectra as ‘Shape-DNA’ of surfaces and solids,” Computer-Aided Design, 2006. DOI: 10.1016/j.cad.2005.10.011.

[17] C. Wachinger, P. Golland, W. Kremen, B. Fischl, M. Reuter, and Alzheimer’s Disease Neuroimaging Initiative, “BrainPrint: A discriminative characterization of brain morphology,” eng, NeuroImage, 2015. DOI: 10.1016/j.neuroimage.2015.01.032.

[18] A. Ortega, P. Frossard, J. Kovacevic, J. M. F. Moura, and P. Van-dergheynst, “Graph Signal Processing: Overview, Challenges, and Applications,” Proceedings of the IEEE, 2018. DOI: 10.1109/JPROC.2018.2820126.6

[19] H. Behjat, N. Leonardi, L. Sornmo, and D. Van De Ville, “Anatomically-adapted graph wavelets for improved group-level fMRI activation mapping,” NeuroImage, 2015.

[20] W. Huang, T. A. Bolton, J. D. Medaglia, D. S. Bassett, A. Ribeiro, and D. Van De Ville, “A Graph Signal Processing Perspective on Functional Brain Imaging,” Proceedings of the IEEE, 2018. DOI: 10.1109/JPROC.2018.2798928.

[21] M. G. Preti and D. Van De Ville, “Decoupling of brain function from structure reveals regional behavioral specialization in humans,” Nature communications, 2019.

[22] K. Glomb, J. Rue Queralt, D. Pascucci, M. Defferrard, S. Tourbier, M. Carboni, M. Rubega, S. Vulliemoz, G. Plomp, and P. Hag-mann, “Connectome spectral analysis to track EEG task dynamics on a subsecond scale,” NeuroImage, 2020. DOI: 10.1016/j.neuroimage.2020.117137.

[23] H. Behjat, A. Tarun, D. Abramian, M. Larsson, and D. V. D. Ville, “Voxel-Wise Brain Graphs From Diffusion MRI: Intrinsic Eigenspace Dimensionality and Application to Functional MRI,” IEEE Open Journal of Engineering in Medicine and Biology, 2023. DOI: 10.1109/OJEMB.2023.3267726.

[24] J. C. Pang, K. M. Aquino, M. Oldehinkel, P. A. Robinson, B. D. Fulcher, M. Breakspear, and A. Fornito, Reply to: Commentary on Pang et al. (2023) Nature, en, 2023. DOI: 10.1101/2023.10.06.560797.

[25] J. Faskowitz, D. Moyer, D. A. Handwerker, J. Gonzalez-Castillo, P. A. Bandettini, S. Jbabdi, and R. Betzel, Commentary on Pang et al. (2023) Nature, en, 2023. DOI: 10.1101/2023.07.20.549785.

[26] H. Behjat and M. Larsson, “Spectral characterization of functional MRI data on voxel-resolution cortical graphs,” in 2020 IEEE 17th International Symposium on Biomedical Imaging (ISBI), IEEE, 2020.

[27] H. Behjat, I. Aganj, D. Abramian, A. Eklund, and C.-F. Westin, “Characterization of spatial dynamics of fMRI data in white matter using diffusion-informed white matter harmonics,” in 2021 IEEE 18th International Symposium on Biomedical Imaging (ISBI), IEEE, 2021.

[28] A. Tarun, H. Behjat, T. Bolton, D. Abramian, and D. Van De Ville, “Structural mediation of human brain activity revealed by white-matter interpolation of fMRI,” Neuroimage, 2020.

[29] D. Abramian, M. Larsson, A. Eklund, I. Aganj, C.-F. Westin, and H. Behjat, “Diffusion-informed spatial smoothing of fMRI data in white matter using spectral graph filters,” Neuroimage, 2021.

